# Neural Representation of The Self

**DOI:** 10.1101/2022.10.19.512950

**Authors:** Marie Levorsen, Ryuta Aoki, Kenji Matsumoto, Constantine Sedikides, Keise Izuma

**Affiliations:** School of Psychology, University of Southampton, Southampton, SO17 1BJ, UK; Graduate School of Humanities, Tokyo Metropolitan University, Tokyo, Japan; Brain Science Institute, Tamagawa University, 6-1-1 Tamagawagakuen, Machida, Tokyo 194-8610, Japan; School of Economics & Management, Kochi University of Technology, Kochi 780-8515, Japan; Research Institute for Future Design, Kochi University of Technology, Kochi 780-8515, Japan

## Abstract

Knowledge about one’s personality, the self-concept, shapes human experience. Social cognitive neuroscience has made strides addressing the question of where and how the self is represented in the brain. The answer, however, remains elusive. We conducted two functional magnetic resonance imaging experiments (with the second being preregistered) employing a self-reference task with a broad range of attributes and carrying out a searchlight representational similarity analysis. The importance of attributes to self-identity was represented in the medial prefrontal cortex (mPFC), whereas mPFC activation was unrelated both to self-descriptiveness of attributes (Experiments 1-2) and importance of attributes to a friend’s self-identity (Experiment 2). Our research provides a comprehensive answer to the abovementioned question: The self-concept is conceptualized in terms of self-importance and represented in the mPFC.

## Introduction

The sense of self shapes human experience^1^. The self (or self-concept) consists of knowledge that people possess about the kind of person they are, such as their names, traits, physical attributes, likes, dislikes beliefs, values, or ingroups (i.e., groups to which they belong) ^2^. The self has been of keen interest to psychologists since the birth of the discipline^3^, and this interest has been rekindled in the last decades^4,5^. From the late 90s^6^ onward, cognitive neuroscientists have been investigating the neural basis of the self via neuroimaging methods^7^. However, where and how the self is represented in the brain remains elusive.

Past neuroimaging studies on the self have typically used a self-reference task^8^ in which individuals are presented with a trait adjective (e.g., honest) and asked to judge whether it describes them. These studies identified a network of brain regions, including medial prefrontal cortex (mPFC) and posterior cingulate cortex (PCC), that are consistently active during the self-reference judgement compared to other semantic (e.g., social desirability) judgements^9–11^.

However, the approach of focusing on differential neural responses during the self-reference versus control tasks to unveil the neural basis of the self has several limitations. First, activation observed by simply comparing the strength of neural responses between the self-reference and control tasks may be due to cognitive processes unrelated to the self. Although the self-reference task is robustly associated with mPFC and PCC activation, the same regions are also active during an other-reference task (e.g., judging if a trait adjective describes one’s best friend) ^9,10,12^, and there are a few cognitive processes common both to self- and other-reference processing^13,14^. For example, autobiographical memory activates the mPFC and PCC, among other regions^15^. Furthermore, positive affect (subjective value) is also linked to the mPFC^16^, and some researchers have argued that the self is related to reward and valuation^17,18^. This limitation is at least partially addressed by recent functional magnetic resonance imaging (fMRI) studies that used a multivariate pattern analysis (MVPA) ^19–24^. In three such studies ^21–23^, mPFC activation patterns were largely separable for self- and other-reference processing. In another study^20^, mPFC activation pattens were more similar for self versus close others than for self versus acquaintances and celebrities. However, although these studies suggested that information processed in the mPFC was different for self-reference versus other-reference, they came short of documenting what information about the self is specifically being processed. Another MVPA study^19^ illustrated that activation patterns in the anterior ventral part of the mPFC (anterior vmPFC) were similar for self-reference processing and positive affect (i.e., viewing positive images; see also ref^24^ for an analogous result), indicating that positive affect is represented in this region during the self-reference task. Taken together, although the recent MVPA literature has advanced understanding of the neural bases of the self, the precise nature of self-referent information that is processed—especially in a more dorsal part of the mPFC—remains unknown.

Second, the majority of previous studies used only trait adjectives as experimental stimuli. This practice likely limits researchers’ ability to identify the neural representations of the self. As stated above, the self includes not only personality traits, but also physical characteristics, preferences, values, achievements, aspirations, abilities, and social groups^25^; thus, personality traits comprise a narrow subset of the self. For example, social identity theory^26,27^ and self-categorization theory^28,29^ advocate the relevance of social or group identity in influencing attitudes, judgements, and behaviors. Using only trait adjectives neglects, to a great degree, this crucial^30^ social aspect of the self. The Twenty Statement Test (TST) ^31^ assesses the content of the self by asking participants to freely provide up to 20 responses to the prompt “I am __ __.” In a review of the self-content literature^32^, among eight studies that used the TST in various countries—including India, Japan, Kenya, and USA—the average proportion of participants’ personality traits was 26.6% (range = 0-58%), whereas that of their social/collective attributes was 28.9% (range = 1-84%). Furthermore, two studies^32,33^ documented that the TST overestimates the proportion of traits in the self-concept. For example, one of those studies^32^ compared the TST with the Writing About Yourself (WAY) task in which participants are provided with a single sheet of blank paper labeled “Writing about Yourself” and asked to write one or more paragraphs about themselves. Across four countries (Australia, Mexico, Philippines, USA), the proportion of personality traits was higher for the TST (39-54%) than the WAY task (9-13%), whereas the proportion of social/collective attributes was similar (21-33%). Taken together, neuroimaging studies that have used traits as experimental stimuli assessed a small part of the self-concept.

Third, most of the neuroimaging research has operationalized the self-concept in terms of trait self-descriptiveness. Although some studies found that mPFC activation increased as a function of trait self-descriptiveness ^22,34–37^, other studies with a similar event-related design did not report results relevant to this association (although they could^38–41^). There is virtually an infinite number of characteristics that can describe an individual, but most of them are irrelevant to self-definition (e.g., “I sleep every day,” “I take the number 38 bus to work,” “I go shopping on weekends”). Stated otherwise, just because an item is self-descriptive does not necessarily mean it is a part of the self-concept^42^. Thus, it is unlikely that the brain has a dedicated neural system for processing information about self-descriptiveness. However, the self might be represented in the brain in terms of personal importance of each characteristic (hereafter, self-importance or centrality). The relevance of taking into account self-importance (as well as self-descriptiveness) when assessing the self has been recognized by psychologists^42,43^. That is, whether information will influence one’s behavior depends on its personal importance. In particular, degree of informational importance may moderate the link between personality and behavior^44^. For example, if being a mother is important to an individual, her behaviours as a mother are likely to be different (e.g., more attentive, careful, responsible, or consistent) from her behaviours in other, less self-defining roles. There is empirical support for this claim^1^. Participants whose moral identity is high (than low) on self-importance are more likely to help outgroup members^45^. Also, how self-important (central) a personality trait is affects how a person seeks^46^ and remembers^47^ information about themselves. Thus, it is likely that there is a dedicated neural system in the brain which encodes the self-importance of incoming information or stimuli^44,48^.

We addressed the question of where and how the self-concept is represented in the brain in two fMRI experiments (the second being preregistered). In doing so, we overcame the abovementioned limitations. Specifically, we used the self-reference task with a broad range of stimuli combined with a representational similarity analysis (RSA) ^49^ of fMRI data to test how mPFC activation patterns are related to self-importance as well as self-descriptiveness. To cover as widely as possible the content of participants’ self-concept, we asked them to complete a task similar to the TST before an fMRI session; our aim was to prepare a stimulus set that was unique to each participant and varied extensively on self-importance and self-descriptiveness. We examined if self-importance and self-descriptiveness of each stimulus are represented in the brain while controlling for other factors (i.e., valence, familiarity, autobiographical memory, stimulus word-length, whether the item was provided by the participant or not). Among several brain regions previously implicated in self-processing^9–11^, we focused especially on the mPFC, as a lesion study indicated that this region is necessary for a stable and accurate self-concept (but is not critical for knowledge about another person) ^50^.

## Methods

### Experiment 1

#### Participants

We recruited 32 right-handed undergraduate students from Tamagawa University. The students had no history of psychiatric disorders. We excluded data from four participants due to excessive head movement (>3mm; 1 participant), because their response consistency in the fMRI tasks was close to chance (1 participant), due to no variance in the post-scan memory rating (1 participant), and because their self-reference rating reliability was low (1 participant). In regard to the last case, each participant completed the self-reference task three times, and we computed correlation coefficient across the three ratings. The average correlation of this fourth participant was 0.21, which was >3 SDs below the group average of *r* = 0.72 [SD = 0.14], suggesting very poor compliance with the task instructions and/or having a highly unstable self-concept. The final sample consisted of 28 participants (16 women, 12 men) aged 19-22 years (*M*_age_ = 19.84, *SD*_age_ = 0.86). Participants ticked a box to indicate their consent prior to the online questionnaires. Additionally, we obtained written consent from all participants prior to the fMRI experiment. The study was approved by both the University of Southampton and Tamagawa University ethics committees. We remunerated each participant with 5,000 Japanese yen.

#### Experimental procedure

The experiment comprised the three following parts, which took place on three separate days: (a) first online questionnaire, (b) second online questionnaire, (c) fMRI experiment. We administered the two online questionnaires in an effort to create the stimulus set for the fMRI experiment—a stimulus set that would be personal and specific to each participant (see below). The first and second online questionnaires were separated by an average of 6.25 days (*SD* = 3.26). The second online questionnaire and fMRI experiment were separated by an average of 8.29 days (*SD* = 2.19).

##### First online questionnaire

During the first online questionnaire, we instructed participants to provide at least 30 characteristics by responding to the prompt “I__”. To facilitate this task, we gave participants examples such as physical characteristics (e.g., I am tall), personality (e.g., I am social), likes or dislikes (e.g., food, music, artists), and groups to which they belonged (university, department, clubs) (Supplemental Information).

##### Second online questionnaire

The second online questionnaire included a total of 80 items some of which were provided by participants during the first online questionnaire, and others were added by an experimenter. Items prepared by the experimenter were intended to dissociate self-descriptiveness, self-importance, and other factors described below. For example, “right-handed” was added with an aim to dissociate self-descriptiveness and self-importance. “School trip” was added with an aim to dissociate self-descriptiveness/self-importance and autobiographical memory. “Convenience store” was added with an aim to dissociate self-descriptiveness/importance and familiarity. We instructed participants to rate each item in terms of (a) self-descriptiveness (1 = *not descriptive at all*, 7 = *very descriptive*), (b) importance to their self-identity (1 = *not important at all*, 7 = *very important*), (c) valence (1 = *very negative*, 7 = *very positive*), (d) familiarity (1= *not familiar at all*, 7 = *very familiar*), and (e) autobiographical memory or the extent to which “each word/phrase brought back memories of your past when you saw it” (1 = *it did not evoke any memory at all*, 7 = *it evoked very vivid memory*).

##### Stimulus set preparation

Based on the ratings, we selected a final stimulus set of 40 items under the stipulation that the ratings not be highly inter-correlated (i.e., effects on neural activities be dissociable). Specifically, we randomly picked 40 out of the 80 items used in the second questionnaire and computed correlations across the following six ratings/characteristics of the randomly selected 40 items: (a) self-descriptiveness, (b) valence, (c) familiarity, (d) autobiographical memory, (e) number of characters, (f) whether the item was provided by a participant during the first questionnaire (1) or not (0). We then recorded the highest correlation coefficient (*r_highest_*). We repeated this process 1,000,000,000 times and selected the final set of 40 items that had the lowest *r_highest_*. For some participants, the self-descriptiveness and self-importance ratings were highly positively correlated. In such a case, we set a different criterion for that correlation. For instance, we computed *r_highest_* without considering *r_self-descriptiveness/self-importance_*, and selected a set of 40 items whose *r_highest_* was the lowest given that *r_self-descriptiveness/self-importance_* was less than 0.6. See Supplemental Information Table 1S for examples of items.

##### fMRI experiment

Prior to the fMRI scan, participants received instructions regarding MRI safety and tasks they would perform inside the fMRI scanner. During the fMRI session, participants performed two tasks: self-reference and word-class judgement (Figure 1). We programmed both of them using Psychtoolbox (http://psychtoolbox.org/) with Matlab software (version 2018a, http://www.mathworks.co.uk). Participants completed six runs of each task. Each run consisted of 40 trials, one item per trial. We presented the same set of 40 items in both tasks and in each run. We counterbalanced task (ABBAABBAABBA or BAABBAABBAAB), and randomized trial order within each run.

**Figure 1.**
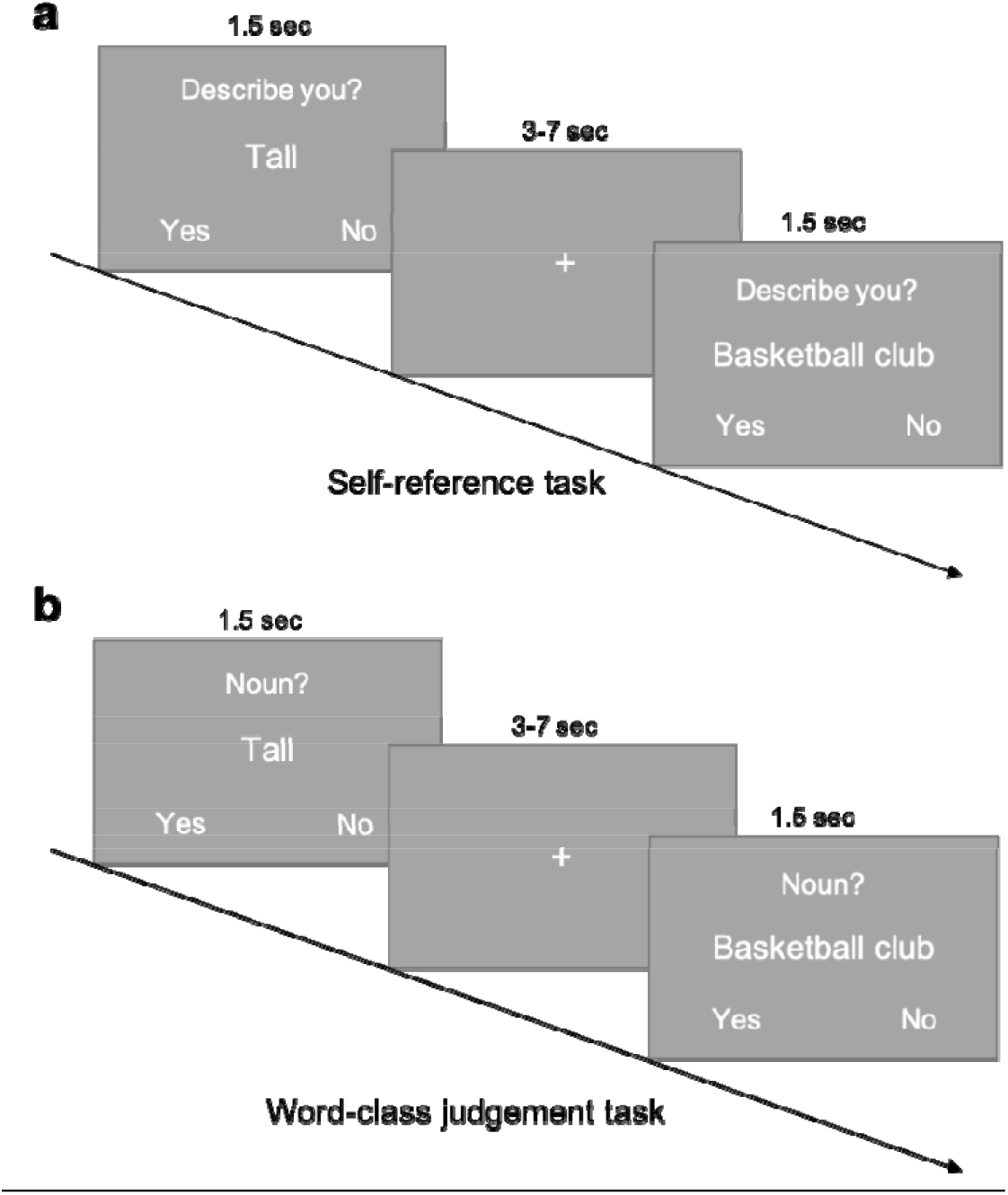
Example of trials during the experimental tasks of self-reference and word-class judgement. (**a**) In the self-reference task, participants viewed characteristics and judged whether each described them. (**b**) In the word-class judgement task, participants viewed characteristics and judged whether each was a noun.

In the self-reference task, for each trial, participants viewed an item. On the screen, above the characteristic, they encountered the question “Describes you?”. For each trial, they could answer “yes” or “no” to indicate whether the characteristic described them or not. For each trial in the word-class judgement task, participants viewed an item. On the screen, above the characteristic, they encountered the question “Noun?”. They could answer “yes” or “no” to indicate whether the characteristic was a noun or not (Figure 1). For both tasks, each item was presented for 1.5 seconds, followed by inter-trial-interval (ITI) (3-7 seconds, mean = 5 seconds). Participants answered by pressing one of two buttons on a response box.

How vividly each item evoked a personal memory might differ between the second questionnaire and the fMRI task. Consequently, after the fMRI scan, we instructed participants to rate the same 40 items on autobiographical memory, namely, to what extent each item evoked an autobiographical memory when seeing it inside the fMRI scanner (1 = *it did not evoke any memory at all*, 7 = *it evoked very vivid memory*). Furthermore, to check for consistency of the self-descriptiveness judgement, we instructed participants to rate the 40 items again on self-descriptiveness using the same response scale. Next, participants completed a demographic questionnaire.

#### Data analysis

##### Behavioral data analysis

Each participant rated each of the 40 items on self-descriptiveness, and they did so three times: (a) during the second online questionnaire, (b) during the fMRI scan, (c) after the fMRI scan. Although participants rated each item 6 times (across 6 fMRI runs) on a 2-point scale (yes or no) during the fMRI scan, they rated each item once on a 7-point scale during the second questionnaire and post-fMRI rating task. Thus, for the self-descriptiveness rating data obtained during the fMRI scan, we computed a self-descriptiveness score for each item as a proportion of yes responses across 6 ratings of each item. We assumed that participants maintained a stable self-concept across a few weeks, and we tested this assumption by checking for consistency of their self-descriptiveness ratings obtained across the three times (or sessions).

#### fMRI data acquisition

We acquired images using a 3-T Trio A Tim MRI (Siemens) scanner with a 32-channel head coil. For functional imaging, we used T2*-weighted gradient-echo echo-planar imaging (EPI) sequences with the following parameters: Time repetition (TR) = 2500 ms, echo time (TE) = 25 ms, flip angle (FA) = 90°, field of view (FOV) = 192 mm^2^, matrix = 64 × 64. We acquired, in an interleaved order, 42 contiguous slices with a thickness of 3 mm. In addition, we acquired a T1-weighted structural image from each participant.

#### fMRI data preprocessing

We carried out preprocessing and statistical analysis of the fMRI data using SPM12 (Welcome Department of Imaging Neuroscience), implemented in MATLAB (Math Works). We discarded the first four volumes before preprocessing and data analyses to allow for T1 equilibration. We conducted preprocessing of the fMRI data with SPM 12’s preproc_fmri.m script starting with realignment of all functional images to a common image. We spatially realigned all images within each run to the first volume of the run using 7th-degree B-spline interpolation, and we unwarped and corrected for motion artefacts. We segmented the T1-weighted structural image and normalized it into a common stereotactic space (MNI atlas). Subsequently, we applied the normalization parameters to the functional images and resampled them to 3 × 3 × 3 mm^3^ isotropic voxels (i.e., original voxel size was retained) using 7th-degree B-spline interpolation. Following the normalization, we spatially smoothed the data (with a Gaussian kernel of 8 mm FWHM) for the univariate analysis. To maintain fine grained activation patterns, we did not apply smoothing prior to the first-level data analysis for the RSA. We applied smoothing before the group analysis of the RSA outputs to account for individual variability in brain structure (with a Gaussian kernel of 4mm FWHM).

#### fMRI data analysis

##### Univariate analysis

We used two general linear models (GLMs) to analyze the fMRI data. In the first GLM, we intended to compare the two conditions (self vs. word), whereas we used the spmT images from the second GLM for the RSA. In the first GLM, we separately modelled 40 self-reference judgement trials and 40 word-class judgement trials using a box-car function convolved with the canonical hemodynamic response function. We submitted the contrast images to a second level analysis. We set the statistical threshold at *p* < 0.005 (uncorrected for multiple comparisons) with a cluster threshold of *p* < 0.05 (Family-Wise Error [FWE] corrected). In the second GLM, we modelled separately each of the 40 items for each task. In the both GLMs, we included six head motion parameters and session effects as nuisance regressors.

##### Representational similarity analysis (RSA): Model representational similarity matrix (RSM)

To test the effect of self-descriptiveness and self-importance on neural responses in the mPFC, while controlling for other factors (i.e., valence, familiarity, autobiographical memory, word-length, whether items were self-provided or not), we used RSA with a searchlight approach. For each participant, we calculated a model RSM separately for the following seven dimensions: (a) self-descriptiveness, (b) self-importance, (c) valence, (d) familiarity, (e) autobiographical memory, (f) word-length, (g) whether each item was self-provided (1) or not (0). For self-descriptiveness, self-importance, valence, and familiarity, we used ratings from the second online questionnaire. For autobiographical memory, we used ratings from the post-scan behavioral session. Each RSM was a 40 × 40 matrix (Figure 2), where each cell represented the similarity of the ratings between two items. For ratings completed on a 7-point scale, we calculated similarity as seven minus the absolute difference between two ratings. For the word length, we calculated similarity as the maximum number of characters in the 40 items minus the absolute difference in the number of characters between two items. Lastly, for whether items were self-provided or not, we coded similarity as 0 if an item was provided by the participant but the other item was not (or *vice versa*), whereas we coded similarity as 1 otherwise (i.e., both items were provided by the participants or both items were provided by the experimenter). We standardized the values with the respective mean and SD for each rating prior to regression analyses.

**Figure 2.**
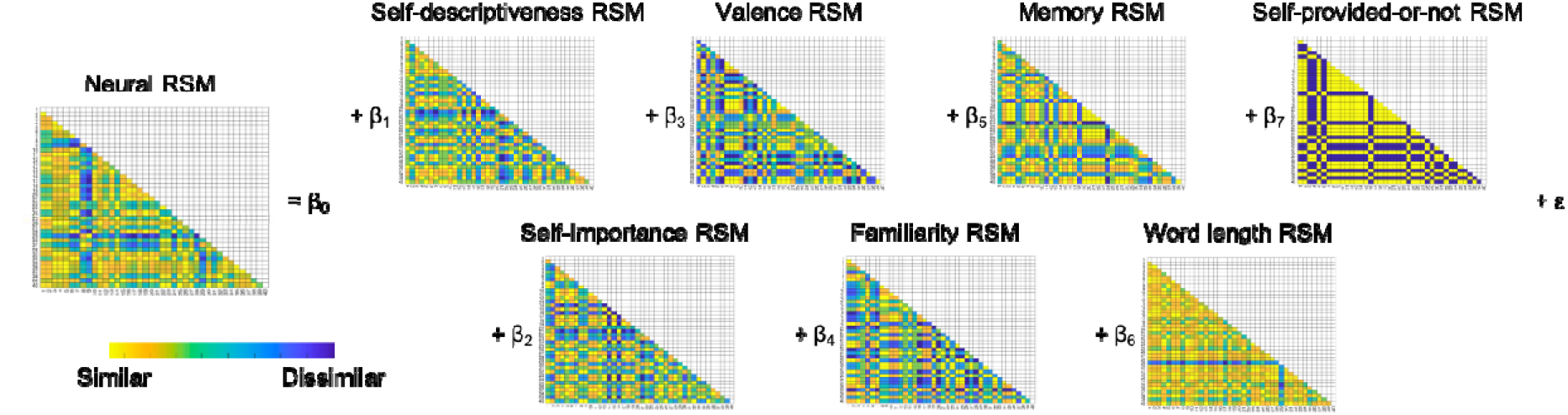
Neural and model RSMs in the multiple regression analyses. For each participant, we made seven model RSMs based on participant ratings and item characteristics. The value in each cell indicates a similarity between a pair of items on a given dimension. We made a neural RSM for each searchlight. We carried out a multiple regression analyses for each searchlight with the seven model RSMs.

##### RSA: NeuralRSM

We extracted local patterns of neural activity from searchlights with a 3-voxel radius, so that each searchlight consisted of a maximum of 123 voxels (and less on the edges of the brain). For each searchlight, we calculated voxel-by-voxel correlations between each pair of the 40 items, which resulted in a 40 × 40 neural RSM (Figure 2). Consequently, the correlation in neural activities between two items within a searchlight was represented by a cell in the respective neural RSM. We Fisher-Z transformed the correlation values prior to further analyses.

##### RSA: Multiple regression analysis

In each searchlight, we conducted a multiple regression analysis, where the seven model RSMs were independent variables and the neural RSM was dependent variable (Figure 2). We repeated this analysis for every searchlight across the mPFC region of interest (ROI; see below for more information about the mPFC mask applied) and the whole brain, resulting in a beta-map for each of the seven independent variables for each of the two tasks (i.e., a total of 14 beta maps for each participant).

##### RSA: Group analysis

We entered the beta-maps into a group-level analysis that we computed with permutation testing (i.e., one-sample t-test with 5,000 permutations) using the Statistical NonParametric Mapping (SnPM) toolbox for SPM^51^. We applied an mPFC mask to the analysis to limit the group-analysis to voxels within the *a priori* ROI. We created the mask with the WFU PickAtlas toolbox for SPM^52^. The mPFC ROI mask included Frontal_Sup_medial_L, Frontal_Sup_medial_R, Frontal_Mid_Orb_L, Frontal_Mid_Orb_R, Rectus_L, Rectus_R, Cingulum_Ant_L, and Cingulum_Ant_R (dilation factor = 2), which we took from the Anatomical Automatic Labeling (AAL) masks implemented in the WFU pickatlas toolbox. We set statistical threshold at *p* < 0.005 with cluster-*p* < 0.05 (FWE corrected).

## Results

### Behavioral results

Participants rated each of the 40 items on self-descriptiveness three times; (a) during the second online questionnaire, (b) during the fMRI scan, (c) after the fMRI scan. Their responses were highly consistent across the three sessions (average within-individual correlation = 0.74; Supplementary Information Table 2S). This finding supports our assumption that participants’ self-concept was stable over the weeks of testing.

We present average correlations between the behavioural ratings (self-descriptiveness, self-importance, valence, familiarity, autobiographical memory, word-length, whether items were self-provided) from the second questionnaire in Figure 3a (note that memory ratings were from the second memory rating task after the fMRI scan). Similarly, we present average correlations across the seven model RSMs in Figure 3b. The correlation between the self-descriptiveness and self-importance model RSMs was the highest (average *r* = 0.31 [*SD* = 0.26]). We also calculated and checked the variance inflation factors (VIFs) for the seven independent variables within each participant. VIF provides an index of the degree to which the variance of a coefficient is increased because of collinearity, with values of above 10 often considered problematic. Across a total of 196 (7 variables × 28 participants) VIFs, 193 of them were below 2, and the maximum VIF was 3.03, indicating reasonable ability to draw inferences on the unique variance explained by each variable in all participants.

**Figure 3.**
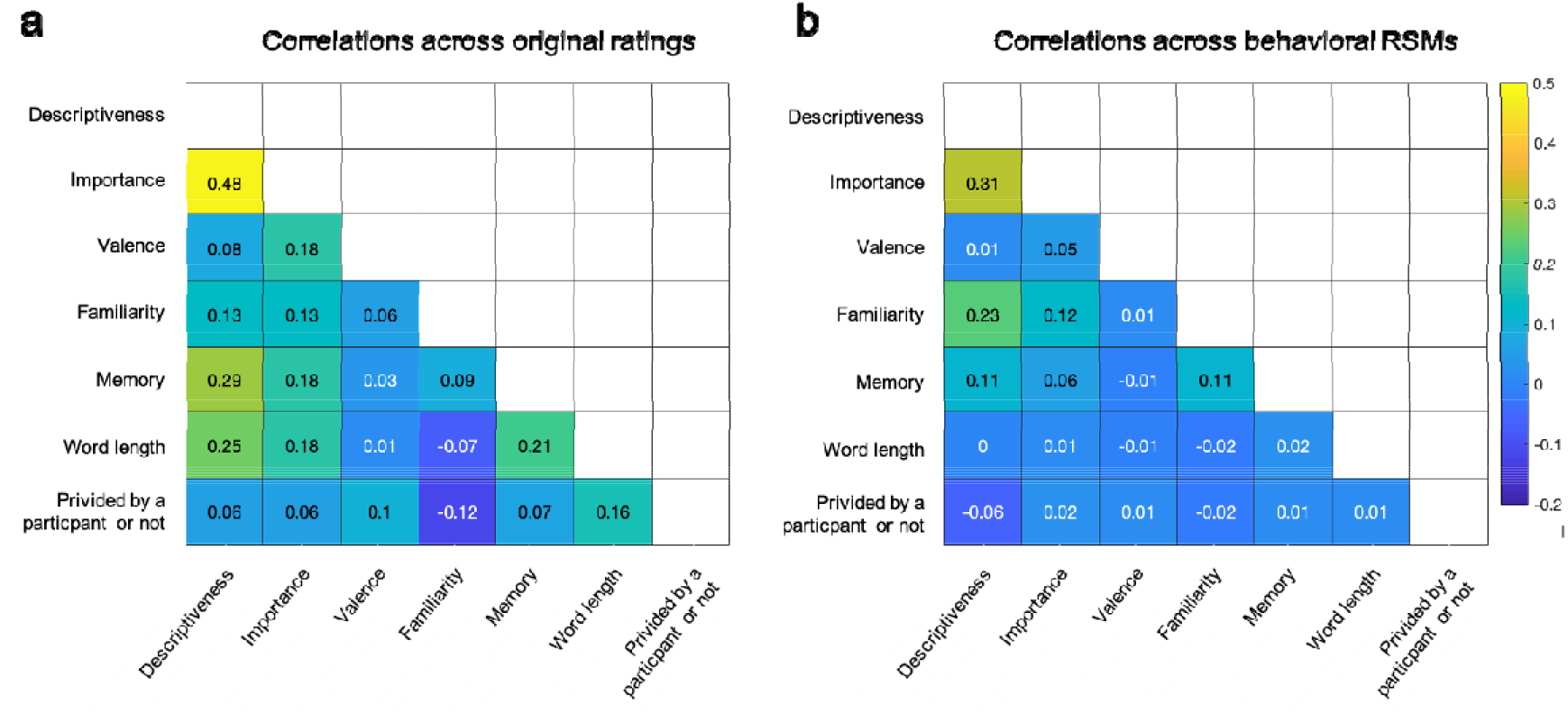
Average correlations across seven ratings (**a**) and seven model RSMs (**b**).

### fMRI results: Univariate analysis

Successfully replicating the previous studies^9–11^, we found that the self-reference versus word-class judgement contrast significantly activated the mPFC and PCC (Supplementary Information Figure 1S). Other activated regions included left and right temporoparietal junction (TPJ), left superior temporal sulcus (STS), and lingual gyrus (Supplementary Information Figure 1S and Supplementary Information Table 3S). The opposite contrast (word vs. self) activated the left inferior frontal gyrus (IFG), which is known to play a major role in language processing^53^ (Supplementary Information Table 3S).

### Searchlight RSA result

We conducted searchlight RSA within the mPFC ROI to test whether self-descriptiveness or self-importance had an effect on the local patterns of activation within the mPFC. We found that different levels of self-importance were represented by different patterns of activation within the mPFC (medial superior frontal gyrus; x = −9, y = 53, z = 29, 306 voxels) during the self-reference task (Figure 4). However, self-descriptiveness was not significantly associated with activation patterns within the mPFC. Likewise, the remaining five variables were not significantly associated with mPFC activations. The mPFC region associated with self-importance (Figure 5a) largely overlapped with the mPFC region activated by the self versus word contrast (Figure 4). Out of the 306 voxels whose activities were significantly associated with self-importance, 194 voxels (63.3%) were included in the area significantly activated by the self-reference task compared to the word task (Figure 5b).

**Figure 4.**
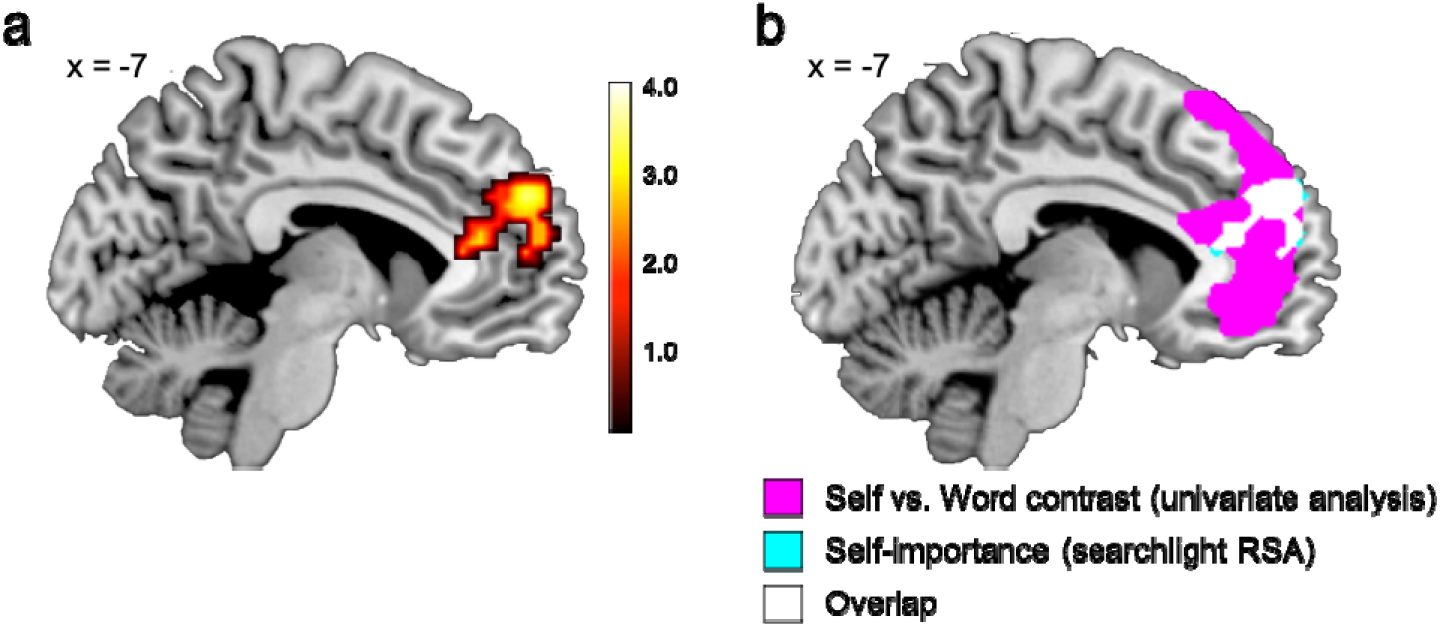
Searchlight RSA result. (**a**) Self-importance was significantly associated with activation patterns within the mPFC during the self-reference task. *p* < 0.005 (uncorrected) and cluster-*p* < 0.05 (FWE corrected) for a display purpose. (**b**) The mPFC areas significantly associated with self-importance largely overlapped with the areas activated by the self-reference task compared to the word class judgement task.

**Figure 5.**
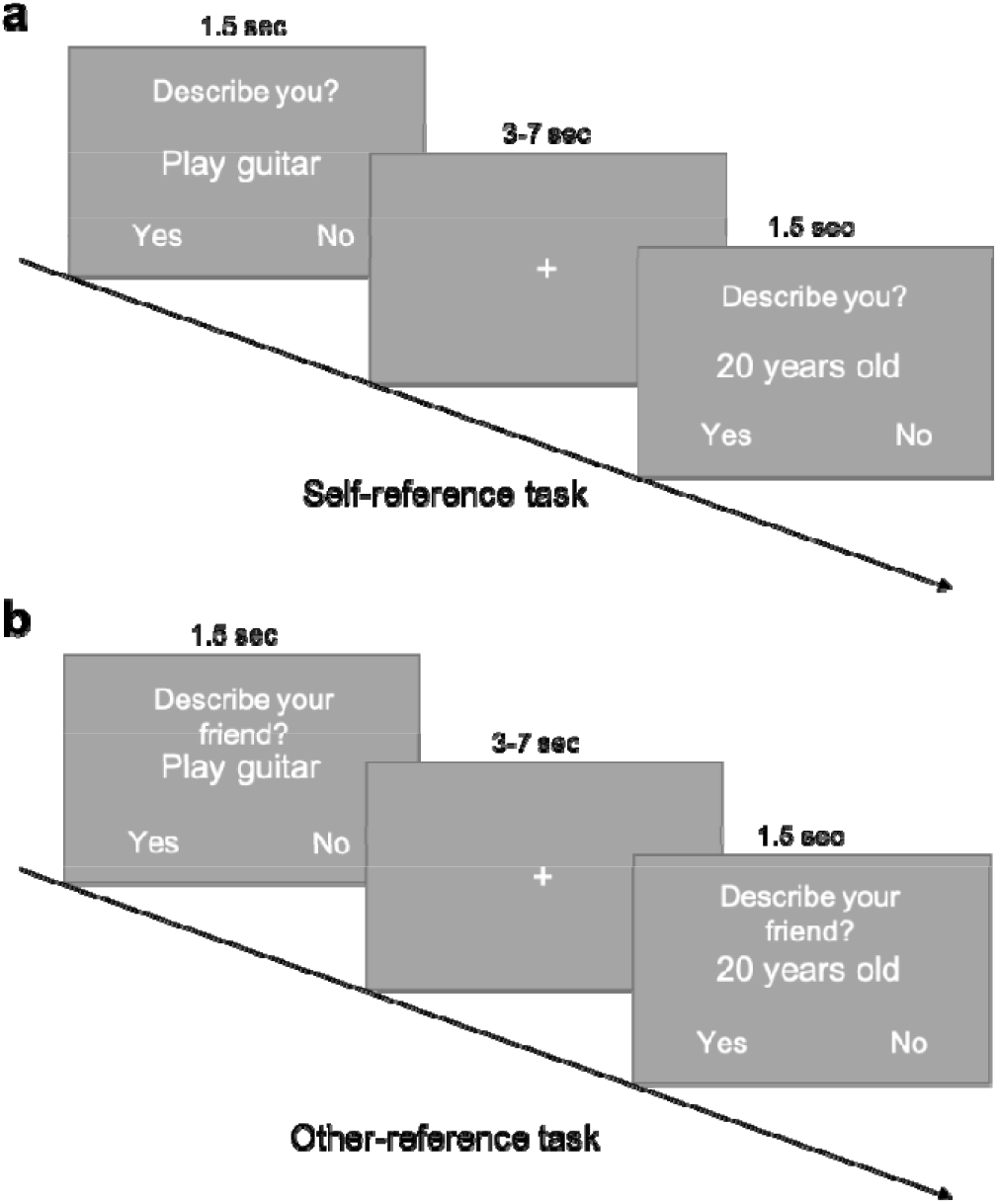
The Self-reference task (**a**) and the Other-reference task (**b**) in Experiment 2.

We repeated the same searchlight RSA using the data from the word class judgement task, and found no significant results. The null effects indicate that the representation of self-importance within the mPFC is task-dependent; instead, self-importance is represented within the mPFC only when performing a task that requires thinking about the self.

Outside of the mPFC ROI, different levels of word-length were represented by different patterns of activation within the visual cortex (lingual-gyrus) for both the word-class judgement task and the self-reference task (Supplementary Information Figure 2S), indicating that visually similar stimuli evoke similar activation patterns in the visual cortex regardless of task. No other significant results emerged.

Taken together, we obtained initial evidence that the mPFC represents self-importance information. However, it is possible that the mPFC represents importance not specific to self-identity, but relevant to another person’s identity as well; that is, the mPFC may not be specific to the self, but instead process person information in general. We addressed this possibility in Experiment 2.

## Experiment 2

### Method

#### Pre-registration

Prior to data analyses, we preregistered the hypotheses, sample size, data analytic plan, and exclusion criteria on the Open Science Framework (https://osf.io/agq3b). We followed the preregistration in all analyses reported below, unless otherwise noted.

#### Participants

As preregistered, our sample comprised 35 undergraduate students (23 men, 12 women) at Kochi University of Technology (*M*_age_ = 19.66, *SD*_age_ = 1.64, range = 18-23 years). All of them were right-handed, and none had a history of psychiatric disorders. We excluded another 8 participants, because they did not meet the preregistration criteria. In particular, we exclude 1 due to a brain anomaly, and 7 because the reliability of either their self-reference or other-reference rating was low. In regard to these 7 participants, we calculated correlations between their responses in the self-reference task and their self-descriptiveness rating in the second questionnaire, as well as the correlation between their other-reference task responses and their friend-descriptiveness rating in the second questionnaire. We considered the correlation value low, if it was < 0.5. All participants ticked a box to indicate their consent prior to the online questionnaires, and they consented in writing prior to the fMRI experiment. The study was approved by the Kochi University of Technology ethics committee. They were remunerated with 2,000 Japanese yen.

#### Power analysis

We carried out a power analysis using the Bootstrap procedure. First, we randomly sampled 35 participants from the 28 participants of Experiment 1, with replacement. For each randomly-selected sample, we conducted a group analysis (one-sample t-test). Given that in Experiment 1 we found significant activations within the mPFC, we applied the same mPFC mask created via the WFU PickAtlas toolbox for SPM^52^. We applied a voxel-wise threshold of *p* < 0.005 (uncorrected) and cluster-*p* < 0.05 (FWE corrected) to assess significance. We repeated these steps 2,000 times, and counted the number of times we found significant activations within the mPFC mask. The result indicated that the sample size of *n* = 35 would achieve power of 91.85%.

#### Experimental procedure

The procedure was similar to that of Experiment 1, consisting of three parts (two online questionnaires, fMRI experiment) on three separate days. The only alteration involved the control task. To examine if the Experiment 1 results were specific to the self, or if the mPFC also encodes important information for a friend’s identity, we used an other-reference task (Figure 5) as control. The first and second online questionnaires were separated by an average of 11.0 days (*SD* = 6.68). The second online questionnaire and fMRI experiment were separated by an average of 14.03 days (*SD* = 8.34).

##### Online questionnaires

As in Experiment 1, in the first online questionnaire, participants provided at least 30 characteristics by responding to the prompt “I __”. Similarly, participants provided the name of a close friend and at least 30 characteristics they believed to be descriptive of or important for that friend. They did so by responding to the prompt “My friend __”.

In the second online questionnaire, participants rated 80 items, some of which were made available by participants during the first online questionnaire. In particular, they rated each item on the following seven dimensions: (a) self-descriptiveness (1 = *not at all descriptive*, 7 = *very descriptive*), (b) importance to self-identity (1 = *not at all important*, 7 = *very important*), (c) friend self-descriptiveness (1 = *not at all descriptive*, 7 = *very descriptive*), (d) importance to friend’s identity (1 = *not at all important*, 7 = *very important*), (e) valence (1 = *very negative*, 7 = very positive), (f) familiarity (1 = *not at all familiar*, 7 = *very familiar*), and (g) autobiographical memory (1 = *it did not evoke any memory at all*, 7 = *it evoked very vivid memory*). We selected a stimulus set of 40 items as in Experiment 1 (see Supplemental Information Table 1S for item examples).

##### fMRI experiment

During the fMRI session, participants carried out the self-reference and other-reference tasks. We used the same set of 40 items for both tasks.

Just like in Experiment 1, during the self-reference task, for each trial, the participants viewed one of the 40 items. On the screen, above the characteristic, they saw the question “Describes you?”. For each trial, they answered “yes” or “no” to indicate whether the characteristic described them (Figure 5a). For each trial in the other-reference task, participants similarly viewed an item on the screen. Above the item, they saw the question “Describes your friend?” and answered “yes” or “no” to indicate whether the characteristic described their friend—the same close friend they mentioned during the first online questionnaire (Figure 5b). For both tasks, each item was presented for 1.5 seconds, followed by inter-trial-interval (ITI) (3-7 seconds, mean = 5 seconds). Participants indicated their answers by pressing one of two buttons on a response box.

##### Post-scan behavioural session

After the scan, participants rated the previously presented words on autobiographical memory again (1 = *it did not evoke any memory at all*, 7 = *it evoked very vivid memory*). Next, they completed a demographic questionnaire.

##### fMRI data acquisition

We acquired images using a Siemens 3.0 T Verio MRI scanner with a 64-channel phased array head coil. For functional imaging, we used T2*-weighted gradient-echo echo-planar imaging (EPI) sequences with the following parameters: Time repetition (TR) = 2500 ms, echo time (TE) = 25 ms, flip angle (FA) = 90°, field of view (FOV) = 192 mm^2^, matrix = 64 × 64. We acquired 42 contiguous slices with a thickness of 3 mm, in an interleaved order. Moreover, we acquired from each participant a high resolution anatomical T1-weighted image (1mm isotropic resolution).

##### fMRI data processing

We conducted preprocessing of fMRI data as in Experiment 1. The preprocessing described in our preregistration stated that we would use an EPI-template when normalizing fMRI data to the standard MNI space. Although, based on visual inspection of normalized images, there was no issue with this method when analyzing the fMRI data from Experiment 1, we noticed that fMRI images normalized with this method were consistently smaller in the anterior-to-posterior and left-to-right dimensions (possibly because of the difference in head-coil between the two experiments; 32 channels in Experiment 1 vs. 64 channels in Experiment 2) (see ref^54^ for a similar case). Accordingly, we decided to use a T1-template when normalizing the fMRI data as implemented in the SPM 12’s preproc_fmri.m script. For the sake of consistency, we re-analyzed fMRI data in Experiment 1 with this new preprocessing steps, as reported above. In Experiment 1, we report the re-analyzed data (note that the two preprocessing steps generated virtually identical results).

##### Univariate fMRI analysis

Similar to Experiment 1, we used two GLMs. In the first GLM, we intended to compare the two conditions (self condition vs. other condition), whereas we used the spmT images from the second GLM for the RSA. We separately modelled 40 self-reference judgement trials and 40 other-reference trials using a box-car function convolved with the canonical hemodynamic response function. We submitted the contrast images to a second level analysis. We set the statistical threshold at *p* < 0.005 (uncorrected for multiple comparisons) with a cluster threshold of *p* < 0.05 (FWE corrected). In the second GLM, we modelled separately each of the 40 items for each task. In both GLMs, we included six head motion parameters and session effects as nuisance regressors.

##### Model RSMs

We conducted searchlight RSA as in Experiment 1. However, in addition to testing the effect of self-descriptiveness and self-importance, we tested the effect of friend-descriptiveness and friend-importance, on neural representations. So, for each participant, we calculated a model RSM separately for each of the following nine dimensions: (a) self-descriptiveness, (b) self-importance, (c) friend-descriptiveness, (d) friend-importance, (e) valence, (f) familiarity, (g) autobiographical memory, (h) word-length, and (i) whether the item was self-provided or not. For self-descriptiveness, self-importance, friend-descriptiveness, friend-importance, valence, and familiarity, we used the ratings from the second questionnaire. For autobiographical memory, we used the ratings from the post-scan behavioural session.

##### Neural RSM

We created a neural RSM for each searchlight as in Experiment I.

##### Multiple regression analysis

In each searchlight, we conducted a multiple regression analysis where the nine model RSMs were independent variables and the neural RSM was the dependent variable. We repeated the analysis for every searchlight across the brain, resulting in a beta-map for each of the nine independent variables and each of the two tasks (a total of 18 [2 × 9] beta maps for each participant). Although not preregistered, we attempted another RSA by adding a model RSM based on participants’ average RT for each item (a total of 10 model RSMs), and this additional RSA produced results virtually identical to those reported below.

##### Group analysis

We conducted the second-level group analysis as in Experiment 1 (i.e., using SnPM). We applied the same statistical threshold (*p* < 0.005 with cluster-*p* < 0.05 [FWE corrected]).

##### Classifier-based MVPA (not pre-registered)

We also conducted a classifier-based MVPA analysis that directly compares the effects of self-importance and friend-importance on mPFC activation. We did so in search for evidence that a neural code for information importance is unique to the self. Specifically, we tested whether, during the other-reference task, the mPFC activation patterns evoked by items high (also middle or low) in self-importance task are distinct from activation patterns evoked by items high (also middle or low) in friend-importance.

First, we carried out another GLM analysis where each item was classified into one of the three categories depending on level of self- (and friend-) importance: high, middle, low. Given that the distribution of ratings was different across participants (e.g., with some frequently providing ratings of 6-7, and others frequently providing ratings of 1-2), we used different criteria for different participants when classifying each item into the three categories so that the three categories included roughly an equal number of items. Of note, within each participant, we used the same criterion for the self and other conditions. Thus, in this GLM, when modelling the fMRI data from the self-reference task, we classified 40 items into three categories based on self-importance ratings: (a) self-importance-high, (b) self-importance-middle, (c) self-importance-low. We modelled separately items in each of the three categories. Similarly, for the other-reference task fMRI data, we classified items into three categories in the same way based on the friend-importance ratings (high, middle, or low), and we modelled separately items in each of the three categories. We included six head motion parameters and session effects as nuisance regressors. We then computed an spmT map for each category per fMRI run resulting in 2 (tasks; self vs. other) × 3 (level of importance) × 6 (runs) spmT images per participants, which we used in the subsequent MVPA.

To define independently a self-importance related mPFC ROI, we used a leave-one-participant-out cross-validation procedure^55^. We re-ran the second-level group analysis (the searchlight RSA group analysis described above) 34 times with a different single participant left out in each. We used each second-level analysis to determine an mPFC ROI for each left-out participant. For each participant, we extracted data from a 3-voxel radius sphere surrounding the peak voxel within the mPFC most strongly associated with self-importance ratings. To ascertain that each participant’s ROI was roughly from the same anatomical sub-region within the mPFC, we searched a peak voxel for each participant within a 30-mm sphere surrounding the peak voxel identified by the group analysis with all 35 participants.

We used a linear support vector machine, which we conducted using Matlab in combination with LIBSVM (https://www.csie.ntu.edu.tw/~cjlin/libsvm/)^56^ with a cost parameter of c = 1 (default). We paired each of the self run and friend run in the order of acquisition, and evaluated classification performances with a leave-one-pair-out cross-validation procedure. Thus, using the spmT images from the five runs of each task, we trained a classifier that discriminates activation patterns between self-importance-high versus friend-importance-high items. Then, using the spmT images from the left-out run of each task, we tested if the classifier could discriminate between self-importance-high versus friend-importance-high items. We repeated the procedure six times so that each run-pair served as the testing set once. We averaged six classification accuracy values for each participant. We conducted the same analysis to test if activation patterns are distinct between items low in self-importance versus items low in friend-importance (also, items middle in self-importance vs. middle in friend-importance). We assessed statistical significance with permutation tests where we performed classifications with scuffled labels 1,000 times to obtain a null distribution. P-values were set at 0.05 (one-tailed) and Bonferroni-corrected for three comparisons.

## Results

### Behavioural results

Participants rated each of the 40 items on self-descriptiveness and friend-descriptiveness twice: (a) during the second online questionnaire, (b) during the fMRI scan. Their responses were highly consistent across the two sessions. Average within-individual correlation for self-descriptiveness ratings was 0.78 (*SD* = 0.09), and average within-subject correlation for friend-descriptiveness was 0.74 (*SD* = 0.09).

We present average correlations between the behavioural ratings (self-descriptiveness, friend’s self-descriptiveness, self-importance, friend’s self-importance, valence, familiarity, autobiographical memory, word-length, whether items self-provided) from the second questionnaire in Figure 6. We checked the VIFs for the nine independent variables within each participant. Results showed that all 315 (9 variables × 35 participants) VIFs were below 2, indicating reasonable ability to make inferences on the unique variance explained by each variable in all participants.

**Figure 6.**
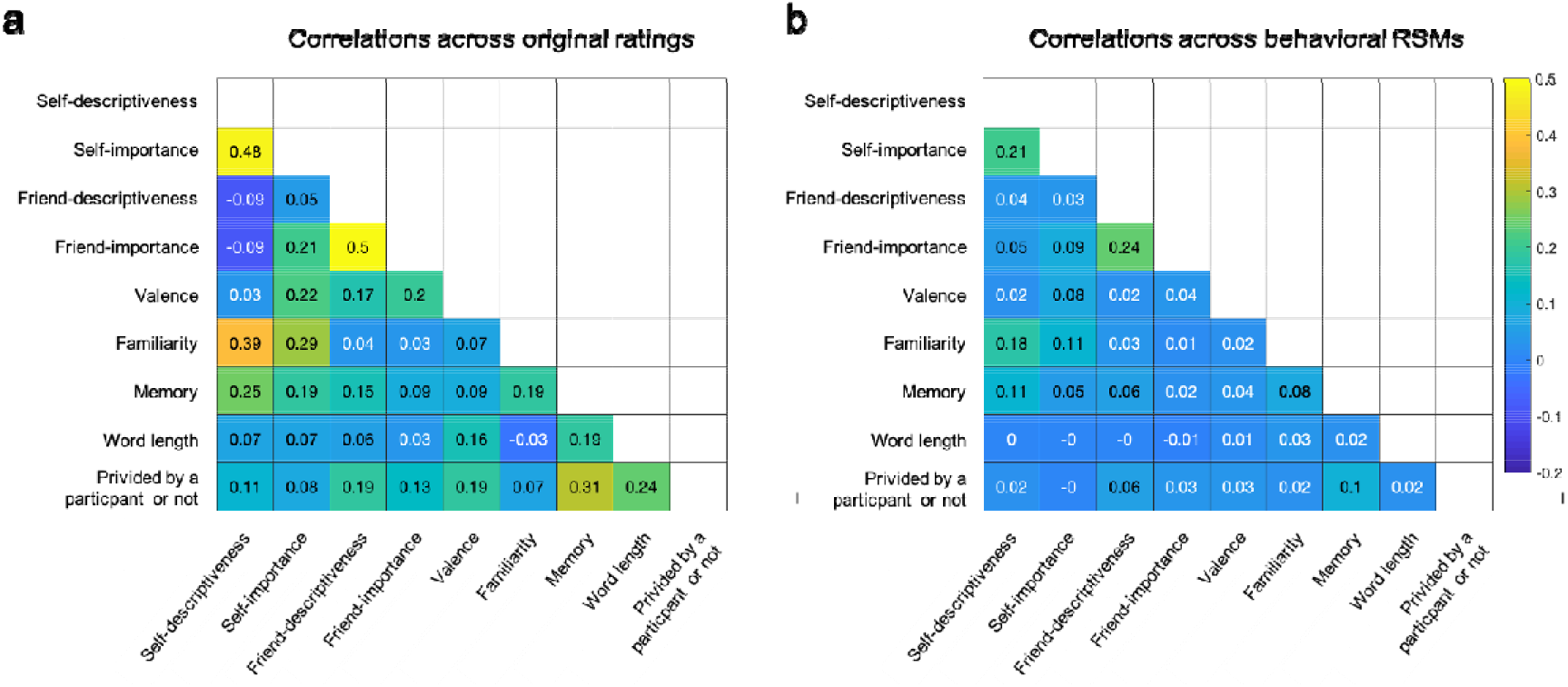
Average correlations across nine ratings (**a**) and nine model RSMs (**b**).

### fMRI results: Univariate analysis

The self-reference versus other reference contrast did not reveal significant activation within the mPFC or across the whole brain. Although this result is in contrast to our preregistered hypothesis, previous studies have generated mixed findings regarding the difference between the self and other conditions, and our finding is consistent with studies that reported no difference (e.g., refs ^40,57–59^). The opposite contrast also did not reveal significant activation in any region.

### Searchlight RSA within the mPFC ROI

Based on our preregistered hypothesis that self-importance is encoded in the mPFC, we first limited the search area to within the mPFC by applying the anatomical mPFC mask. We conducted searchlight RSA to test whether self-importance information is represented in areas within the mPFC during self-reference task. As hypothesized, self-importance was reliably signaled in the mPFC during the self-reference task (x = 3, y = 41, z = 50; 280 voxels; Figure 7). In contrast, self-descriptiveness was not encoded in the mPFC during the self-reference task. Furthermore, neither self-importance nor self-descriptiveness were encoded in the mPFC during the other-reference task. These results replicate those of Experiment 1. Processing of information about how important each stimulus is to the self in the mPFC is task-dependent, and its neural representations emerge only when performing a task that requires thinking about the self.

**Figure 7.**
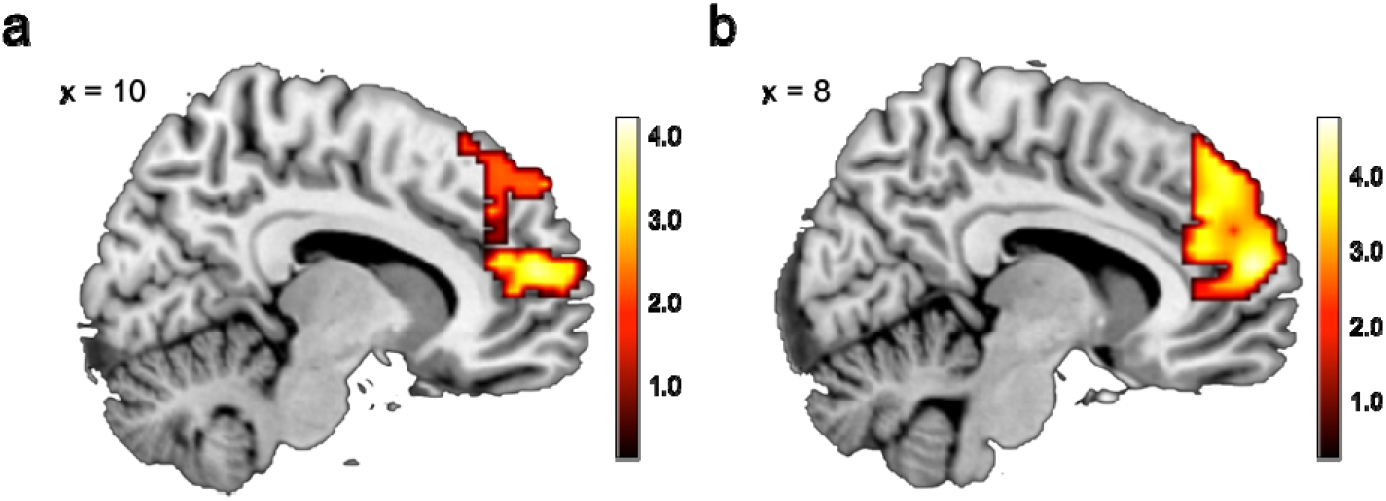
Searchlight RSA result (**a**) and mega-analysis results (**b**). (**a**) Replicating the Experiment 1 findings, self-importance was significantly associated with activation patterns within the mPFC in Experiment 2. *p* < 0.005 (uncorrected) and cluster-*p* < 0.05 (FWE corrected) for display purposes. (**b**) Mega-analysis results (*n* = 63) showing an mPFC cluster that encodes self-importance information. *p* < 0.005 (uncorrected) and cluster-*p* < 0.05 (FWE corrected) for display purposes.

In contrast, both friend-descriptiveness and friend-importance were not significantly associated with mPFC activation during the self-reference and other-reference tasks. Likewise, the remaining five variables were not significantly related to mPFC activation during either task.

### Whole-brain searchlight RSA

We performed searchlight RSA throughout the whole-brain. Consistent with Experiment 1, we found that different levels of word-length were represented by different patterns of activation within the visual cortex (lingual-gyrus) for both the self-reference and other-reference tasks (Supplementary Information Figure 3S). We also found that, during the self-reference task, familiarity ratings were related to activation patterns in left middle frontal gyrus (MFG; x = −21, y = 4, z = 47, 648 voxels) and in left inferior frontal gyrus (IFG; x = −45, y = 11, z = 14, 302 voxels). No other significant effects emerged.

### Exploratory analysis directly comparing effects of self-importance and friend-importance

Although classification performance for items high in importance was not significant, the classifier-based MVPA successfully discriminated activation patterns between self-importance-middle versus friend-importance-middle and between self-importance-low versus friend-importance-low (Table 1), indicating that neural codes_for information importance are largely unique to the self. Classification performance was not significant for items high in importance; however, our additional analysis found that the data were noisier for items high in importance (i.e., activation patterns evoked by the same item were less consistent across six runs of the same task; Supplemental Information Figure 4). It also found that the self-reference task data were noisier compared to those of the other-reference task. Thus, mPFC activation patterns evoked by items high in self-importance during the self-reference task were least consistent (i.e., noisiest) across six runs. This might be because performing the self-reference (vs. other-reference) task and items high (vs. low) in importance evokes other cognitive/affective processes (e.g., autobiographical memory, positive affect) that can influence mPFC activation^15,16^, and these unrelated processes might have impacted on mPFC activation patterns differently in each run. Regardless, the elevated level of noise observed in items high in self-importance explains, at least partially, the non-significance classification performance for items high in self-versus friend-importance.

**Table 1:** Classifier-based MVPA results.

| Comparison | Classification Performance | $P_{perm}$<br>(uncorrected) |
| --- | --- | --- |
| High importance | 50.48% | 0.40 |
| Middle importance | 62.62% | < 0.001** |
| Low importance | 55.71% | 0.003* |

\* $p < 0.05$ and \*\* $p < 0.01$ (Bonferroni corrected). Significant classification performance means that activation patterns were distinct in the self versus other conditions for a given importance level.

### Mega-analysis (notpre-registered)

Given that the self condition was common across Experiments 1 and 2, we combined the self-task data from both experiments and ran a mega-analysis that included 63 participants. Self-importance information was reliably encoded in a large cluster in the mPFC (x = 15, y = 38, z = 26, 942 voxels; Figure 7b).

When we did not apply the anatomical mPFC mask, the above mPFC cluster extended laterally to right IFG (x = 36, y = 38, z = 5) and left IFG (x = −33, y = 29, z = 29) consisting of 2,087 voxels. We observed no other significant cluster for self-importance. Familiarity information was represented in right superior frontal gyrus (SFG; x = 27, y = 26, z = 53; 303 voxels), and autobiographical memory information was represented in right lingual gyrus (x = 27, y = −67, z= −1; 440 voxels). Furthermore, unsurprisingly, word length was strongly associated with activation patterns in the visual cortex (right: x = 12, y = −85, z = 2, and left: x = −9, y = −91, z = 2; a total of 2,413 voxels). We observed no significant result for self-descriptiveness and valence.

## General Discussion

Previous research has linked the self-reference task to neural activation in the mPFC^8^, but it is unclear which information about the self is represented in that brain region. Across two self-reference experiments, while controlling for potential confounds (valence, familiarity, autobiographical memory, word-length, whether items were self-provided), we consistently demonstrated that the mPFC represents how important attributes are to one’s self-identity. The results suggest that the self-concept is represented in the mPFC and conceptualized in terms of self-importance, not self-descriptiveness. In Experiment 2, we did not observe the relationship between mPFC neural responses and importance in the other (best friend) condition, and mPFC activation patterns associated with each of three levels of importance were generally distinct between self and best friend, suggesting that the mPFC represents information about the importance of information specifically to the self. Taken together, our research provides a compelling answer to a long-standing question about how and where the self is represented in the brain.

Although neuroimaging research has documented the involvement of the default mode network in the self-reference task^11,60^, the current results indicate that only the mPFC is associated with self-importance. This finding is consistent with a previous lesion study, which illustrated the mPFC’s crucial role in accurate and reliable trait knowledge of the self^50^. A patient (J.S., 74 years-old white male) had extensive damage to the medial prefrontal areas including orbitofrontal cortex and anterior cingulate gyrus (also extending to anterior and inferior dorsolateral prefrontal areas). He and control participants completed a self-reference task on two occasions (separated by 7-10 days) using the same trait adjectives. A male nurse who had known Patient J.S. for 5 years also rated patient J.S. on the same traits. Patient J.S.’s ratings were less consistent across two sessions and less consistent with ratings done by the nurse compared to the control group (each control participant was also rated by another person who had known them well). When patient J.S. was asked to rate the nurse, his ratings were consistent across two sessions and consistent with ratings done by the nurse himself (no significant difference from the control group was observed), indicating that trait knowledge of another person was preserved. To the best of our knowledge, damage to the mPFC is the only case where trait knowledge of self is impaired, which is in a sharp contrast to other non-self-related knowledge that is impaired after damage to parietal, temporal, or frontal areas^61–63^. In similar experiments with various patients (e.g., Autism, ADHD, Alzheimer’s disease), trait self-knowledge was remarkably resistant to neural and cognitive damage^64^. An example is patient D.B., who suffered profound amnesia due to anoxia following cardiac arrest^65^. Despite his significant memory impairment, patient D.B. exhibited normal (accurate and consistent) trait self-knowledge, whereas trait knowledge of his daughter was significantly less accurate and less consistent compared to control participants. A more recent lesion study also indicated that consistent trait self-knowledge was preserved after damage to anterior ventral temporal lobe, whereas other non-personal semantic knowledge was impaired (especially if the damage was on the left hemisphere) ^66^. In all, the mPFC is the only brain region thus far known to play a crucial role in a consistent and accurate self-concept.

We demonstrated across two experiments that self-importance is represented in the mPFC only during the self-reference task, whereas stimulus’ perceptual properties (i.e., word length) are represented in the visual cortex regardless of the task involved. This task-specific neural representation is consistent with RSA studies on object representations^67^, which showed that information relevant to a given task is represented in prefrontal and parietal areas only while performing the task, whereas occipitotemporal areas mainly represent stimulus’ perceptual properties (e.g., object shape) regardless of task involved. However, during the self-reference task, participants judged whether each personality trait describes them or not; this task does not explicitly require judging how important each trait is to one’s identity. Hence, our results suggest an interesting possibility: Individuals may actively use self-importance information of a stimulus when judging self-descriptiveness. Furthermore, given the lesion study described above^50^, the use of self-importance information represented in the mPFC might be necessary to perform a self-reference task accurately and consistently.

Our findings have far reaching implications. First, they can be contextualized in psychological models of the self-concept. One family of such models depicts the self-concept as an associative network structure where the self is a central entity (node) connected to a number of self-relevant features (e.g., “young,” “university student,” “tall”) that are themselves connected to each other^43,68^. Researchers^69^ further added associative strength to the network model so that some of features (nodes) are more or less strongly connected to the self (and each other). Although these reserachers^69^ considered the strength of association as “the potential for one concept to activate another” (p. 5), its psychological meaning was unspecified. Our findings suggest that, for links (edges) directly connected to the self, strength of association may be understood as degree of self-importance. Given that associative strength is considered responsible for reaction time facilitation or inhibition during the Implicit Association Test (IAT) ^69,70^, our findings generate a hypothesis about reaction time facilitation (e.g., priming effects) based on information self-importance, but not self-descriptiveness (although factors other than self-importance are likely to affect reaction times, such as valence). For example, if being a writer is important to an individual, processing speed for the word “writer” will be facilitated after seeing a prime word “self” (or other highly self-important stimulus). Thus, scores on the self-esteem IAT^71^ might reflect the self-importance information processing function of the mPFC as well as the valence processing function of the reward related network. Largely consistent with this possibility, individual difference in implicit self-esteem as measured by the IAT are independently predicted by activation patterns in the mPFC and those in reward-related brain regions ^72^.

Second, the findings have implications for psychological research on the link between the self-concept and mental health. For example, it possible that people who have greater self-complexity (i.e., higher number of, and great differentiation between, self-aspects) are less likely to experience depression, physical illness, and stress in response to aversive events^73^, especially when they perceive high control over their self-aspects^74^. Similarly, individuals who identify with multiple groups, compared to a single group, report lower stress levels^75^. Furthermore, having multiple group identities after a life transition is associated with better health^76^, and group membership during a life transition is related to enhanced well-being^77^. Other lines of research point to a link between mental conditions or disabilities and the self-concept. For instance, schizophrenia is associated with changes in self-identity^78^, and individuals with autism manifest atypical neural self-representation^79^. How information self-importance is represented in these patients’ brains might shed new light on the nature of mental health, including schizophrenia and autism.

Third, the findings have implications for developmental psychology. The self-concept changes profoundly, especially during adolescence^80^. In a longitudinal fMRI, researchers^81^ tested how brain activations during the self-reference task change from early to mid-adolescents by scanning adolescent participants’ brains once a year over a three-year time period. Activation in the dmPFC during both the self-reference and other-reference tasks increased from the first measurement (*M*_age_ = 12.9) to the second measurement (*M*_age_ = 13.9), but did not differ between the second and last measurement (*M*_age_ = 15.0). Furthermore, activation during the third measurement did not differ from that of young adults. The researchers tentatively proposed that, around age 14, activation in the mPFC during self-evaluations may stabilize and a consistent self-concept be established. Follow-up work should use a combination of multivariate fMRI analysis and longitudinal designs to further test how self-importance representations in the mPFC change during development, and how neural self-representations relate to self-concept stability and possibly to maladaptive behaviour in adolescence^82^.

Fourth, the findings have implications for the long-debated nature of the self among psychologists. One stream of research has emphasized the cognitive properties of the self^83,84^, characterizing its cognitive structure as complex but ordinary^85^. Another stream of research has emphasized the motivational properties of the self ^86,87^, emphasizing its uniqueness ^88,89^. Our findings align with the second empirical stream. If the cognitive representation of the self is unique compared to the cognitive representation of other, this uniqueness lies in motivation (here: attribute self-importance) rather than cognition (here: attribute self-descriptiveness). Moreover, the findings have implications for the long-debated nature of the self among philosophers. Numerous philosophers have cast serious doubts on the mere existence of the self ^90–93^. Here, we countered his viewpoint by providing evidence for the representation of the self in the brain, not only in terms self-descriptiveness (as many other neuroscientists have done), but also in terms of self-importance.

Our findings open up several avenues for research. As stated earlier, the self-concept influences attitudes and behaviours. In a self-domain the individual considers important (e.g., being a parent), their behaviour is more consistent and less susceptible to social influence. Although research has addressed neural mechanisms underlying decision-making and social influence^94–97^, the majority of the literature used stimuli that were unlikely to be important to one’s identity (e.g., faces of strangers, stationaries, snacks; see ref^98^ for an exception), but how people make decisions depends on how important the decision is to their self-identity. Thus, it is worth testing whether (and how) self-representation in the mPFC plays a role in regulating activities in regions implicated in decision-making depending on the importance of a decision to the individual. Specifically, our RSA results suggest that there are populations of neurons in the mPFC, each of which is differentially tuned to the level of self-importance of a presented stimulus. These neural populations might be connected in diverse ways to a network of brain regions implicated in decision-making.

In conclusion, research on the self has a long history in psychology^3^, and the question of “where is the self in the brain?” has attracted keen theoretical and empirical interest in the last two decades^6^. Earlier neuroimaging studies reported a robust link between mPFC activation and self-reference processing^9–11^. More recently, several MVPA studies advanced understanding of the neural representations of the self^19–23,60,99^. However, what information about the self is processed in the mPFC during a self-reference task has eluded an answer. Our research pinned down the nature of the information about the self that is represented in the mPFC: Across two experiments, we demonstrated that information about self-importance (how important a stimulus is to one’s self-identity), but not self-descriptiveness, is represented in the mPFC. Put otherwise, the self-concept is represented in the mPFC in terms of self-importance. The mPFC is a neural locus of the self-concept, and this neural system may play a pivotal role in maintaining a consistent and accurate self-concept.

## Data Availability

Unthresholded group-level statistical maps are available on NeuroVault (https://neurovault.org/collections/13069/)

## Acknowledgements

We thank Atsushi Miyazaki, Mika Oosawa and Maoko Yamanaka for their assistance with fMRI data collection. This research was supported by a Japan Society for Promotion of Science (JSPS) KAKENHI Grant Number JP19K24680 (to K.I.) and the Ministry of Education, Culture, Sports, Science and Technology (MEXT) as part of Joint Research Program implemented at Tamagawa University Brain Science Institute, in Japan.

## Notes

### Competing Interest Statement

The authors have declared no competing interest.

