## Supplemental Information for "Neural Representation of The Self"

#### Instructions

At the beginning of the first online questionnaire, participants received the following instructions:

--------------------------------------------------------------------------------------------

In this first questionnaire, we will ask you who you are.

Please feel free to answer in the format "I ________________."

Please feel free to write about anything that applies to you, such as your physical characteristics (e.g., I am tall), your personality (e.g., I am social), what you like (food, music, artists, etc.; e.g., I like xx) or dislike (I dislike xx), groups you belong to (university, department, clubs, etc.; e.g., I belong to xx clubs), your background (e.g., I graduated from xx High School), values/ideas that you hold dear (e.g., I am an animal rights activist), and so on.

Please tell us not only about your likes and strengths, but also about your dislikes, weaknesses, inhibitions you might have, and other negative things.

You may start a sentence with "I" or "My" (e.g., my favorite word is "xx," "my birthday is on March xx," "my dog's name is xx," etc.).

However, please do not include any information that completely identifies you (e.g., name, e-mail address, etc.). Otherwise, be as specific as possible (e.g., name of high school, clubs you belong to, etc.).

Your answers will be associated with a random ID number and will be stored separately from your name, e-mail address, and other personally identifiable data. However, you do not have to give us anything that you do not want others to know. Please only describe things that you are willing to share.

Some of the words and phrases that you give will be used in a second online questionnaire and fMRI experiment at a later date. The information you provide here will never be used for any purpose other than the experiment.

Please give a minimum of 30 phrases (maximum 40).

When you are ready, please fill in and say a sentence starting with "I" in the answer box below.

--------------------------------------------------------------------------------------------

### Figures

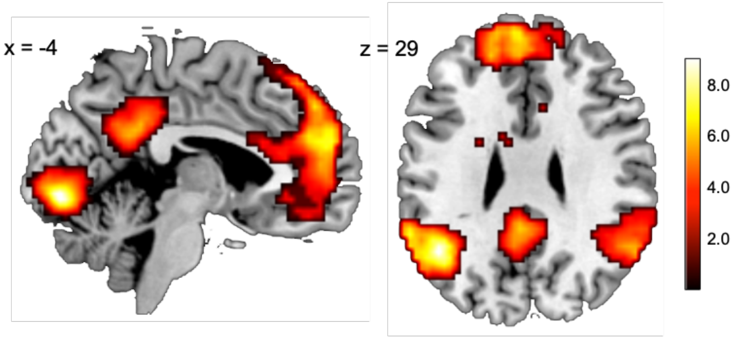


**Figure 1S.** Group activation map for the self versus word contrast. We set up the voxel-wise threshold at $p < 0.005$ (uncorrected), and set up cluster size threshold at $p < 0.05$ (FWE corrected). See Supplementary Information Table 3S for all activated areas.

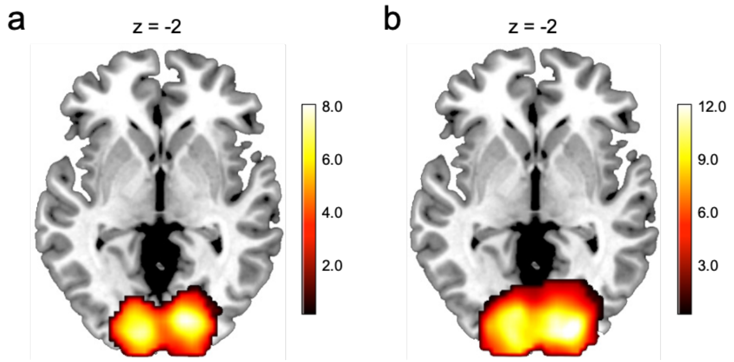


**Figure 2S.** RSA results of the word-length model RSM during the self-reference task (**a**) and word-class judgment task (**b**) in Experiment 1. Activations patterns in the visual cortex were significantly associated with the number of characters in text stimuli for both tasks (Self-reference task; x = 18, y = -82, z = -7, 1,783 voxels; Word-class judgement task; x = 21, y = -88, z = -4, 3,027 voxels). The voxel-wise threshold was set at $p < 0.005$ (uncorrected), and cluster size threshold was set at $p < 0.05$ (FWE corrected).

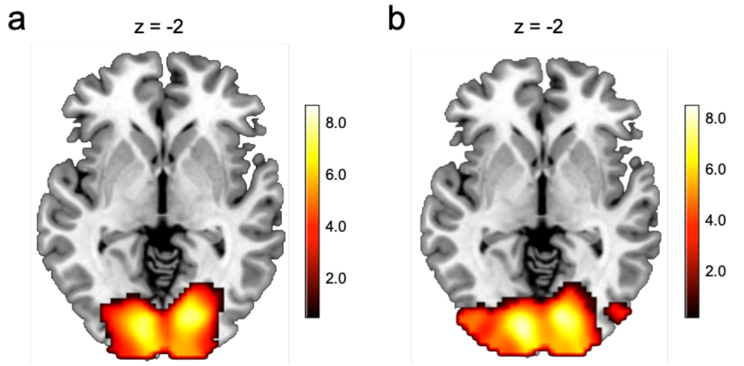


**Figure 3S.** RSA results of the word-length model RSM during the self-reference task (**a**) and other-reference task (**b**) in Experiment 2. Activations patterns in the visual cortex were significantly associated with the number of characters in text stimuli for both tasks (Self-reference task; x = 12, y = -82, z = 2, 2,382 voxels; Friend-reference task; x = 15, y = -88, z = 5, 3,059 voxels). The voxel-wise threshold was set at $p < 0.005$ (uncorrected), and cluster size threshold was set at $p < 0.05$ (FWE corrected).

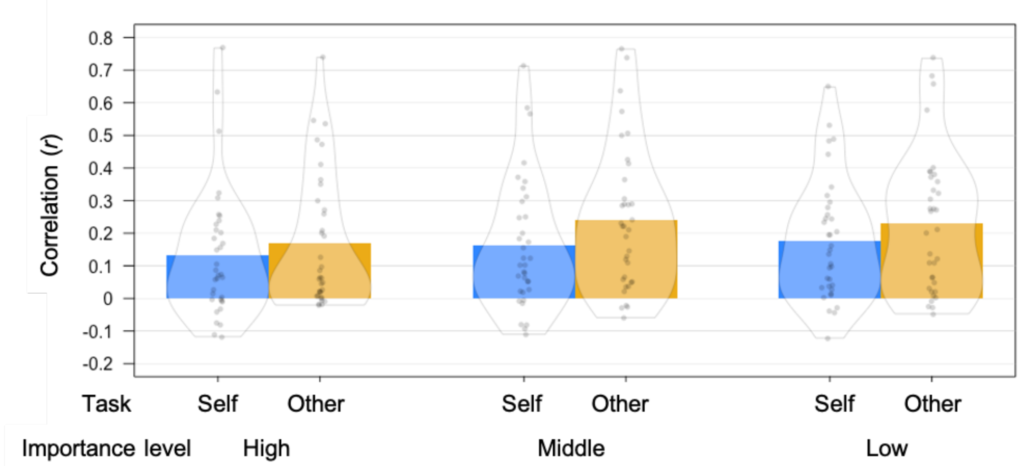


**Figure 4S.** Average within-condition correlations for each of the six conditions (2 [task; self vs. other] × 3 [importance-level; high, middle, or low]). Each participants completed six runs of each task, and within-condition correlations were computed for all possible run-pairs (a total of 15) which was averaged within each participant. A 2 × 3 repeated-measures Analysis of Variance (ANOVA) revealed significant main effects of both task ($F_{(1, 34)} = 6.71$, $p$ = 0.014) and importance-level ($F_{(2, 68)} = 3.67$, $p = 0.0307$), while a task × importance-level interaction was not significant ($F_{(2, 68)} = 0.5262$ $p = 0.539$). Note that the correlation coefficients were Fisher-z transformed before conducting the ANOVA.

### Supplementary Tables

**Table 1S:** Examples of Items Used in the Experiments

| *Physical* | *Social* | *Attributes* | *Other* |
|---|---|---|---|
| Female | Art club | Talkative | Film |
| Bad eyesight | Flower arrangement club | Smart | Twitter |
| Tall | Department of Engineering | Hate prawns | Christmas |
| Hay fever | XX High School graduate | Open-minded | School trip |
| Born in Tokyo | Female | Compassionate | Rain |
| Sweaty | Softball team | Good singer | Piano |
| Brown hair | Japanese | Dog person | Earthquake |
| Sensitive skin | Basketball club | Play guitar | |
| Right-handed | Buddhist | Family-oriented | |
| 20 years old | Kochi University of Technology student | Like to discuss ideas | |

We used 959 unique items across the two experiments. Items in the "Other" category were mainly prepared by an experimenter. The coding scheme is based on Cousins (1989)[1].

**Table 2S:** Average Within-Person Correlations (SD) Across the Three Self-Descriptiveness Ratings in Experiment 1

| | Second questionnaire | fMRI task | Post-fMRI rating |
|---|---|---|---|
| Second questionnaire | - | | |
| fMRI task | 0.83 (0.09)*** | - | |
| Post-fMRI rating | 0.72 (0.14)*** | 0.67 (0.12)*** | - |

Each participant rated each of the 40 items on self-descriptiveness three times; 1) during the second online questionnaire, 2) during the fMRI scan, and 3) after the fMRI scan. *** $p < 0.001$ (corrected for multiple comparisons) based on one sample t-test (one-tailed; correlation coefficients were Fisher-z transformed before the t-tests).

**Table 3S:** Brain Regions Showing Significant Activations During the Self-Reference Task and the Word Class Judgment Task

| Contrast | Location | MNI coordinates | | | Z | Cluster size (voxels) |
|---|---|---|---|---|---|---|
| | | x | y | z | | |
| Self > Word | dmPFC | -9 | 41 | 53 | 5.62 | 1,605 |
| | *mPFC* | -3 | 50 | 23 | 5.42 | |
| | *dACC* | 6 | 20 | 20 | 4.19 | |
| | MFG | -33 | 20 | 38 | 4.68 | 171 |
| | left STS | -60 | -22 | -10 | 5.55 | 736 |
| | PCC | -3 | -46 | 29 | 4.82 | 370 |
| | left TPJ | -45 | -58 | 29 | 6.20 | 580 |
| | right TPJ | 57 | -58 | 29 | 5.83 | 339 |
| | Lingual gyrus | -3 | -85 | -4 | 6.37 | 722 |
| Word > Self | left IFG | -48 | 32 | 20 | 4.69 | 519 |

The statistical threshold was set at $p < 0.005$ (uncorrected for multiple comparisons) with a cluster threshold $p < 0.05$ (FWE corrected). dmPFC; dorsomedial prefrontal cortex, dACC; dorsal anterior cingulate cortex, MFG; middle frontal gyrus, STS; superior temporal sulcus, PCC; posterior cingulate cortex, TPJ; temporoparietal junction, IFG; inferior frontal gyrus. Voxel size = 3 × 3 × 3 mm.
